# Spatial metabolomics using imaging mass spectrometry to identify the localization of asparaptine in *Asparagus officinalis*

**DOI:** 10.1101/2021.01.15.426840

**Authors:** Ryo Nakabayashi, Kei Hashimoto, Tetsuya Mori, Kiminori Toyooka, Hiroshi Sudo, Kazuki Saito

## Abstract

Spatial metabolomics uses imaging mass spectrometry (IMS) to localize metabolites within tissue section. Here, we performed matrix-assisted laser desorption/ionization-Fourier transform ion cyclotron resonance-IMS (MALDI-FTICR-IMS) to identify the localization of asparaptine, a naturally occurring inhibitor of angiotensin-converting enzyme, in green spears of asparagus (*Asparagus officinalis*). Spatial metabolome data were acquired with an untargeted manner. Segmentation analysis using the data characterized tissue-type-dependent and - independent distribution patterns in cross-sections of asparagus spears. Moreover, asparaptine accumulated at high levels in developing lateral shoot tissues. Quantification of asparaptine in lateral shoots using liquid chromatography-tandem mass spectrometry (LC-MS/MS) validated the IMS analysis. These results provide valuable information for understanding the function of asparaptine in asparagus, and identify the lateral shoot as a potential region of interest for multiomic studies to examine gene-to-metabolite associations in asparaptine biosynthesis.

## Introduction

Spatial metabolomics is the study of metabolite localization within tissues, identifying the specific regional pattern of high and low abundance for each metabolite (Boughton et al. 2016; Geier et al. 2020). The functions of many specialized metabolites are largely unknown. Spatial metabolome data may be the key to understanding the function of these metabolites (Dong et al. 2020). By understanding the spatial relationships between metabolites within plant tissue, and combining this knowledge with spatial transcriptome data, it is possible to characterize regional gene-to-metabolite associations and identify candidate biosynthetic genes involved in specialized metabolic pathways. The local function of a metabolite can then be ascertained by comparative analysis of plants altered to lack that metabolite by gene knock-out using genome-editing technologies.

Matrix-assisted laser desorption/ionization-Fourier transform ion cyclotron resonance-imaging mass spectrometry (MALDI-FTICR-IMS) is an analytical method to visualize site-specific metabolite signal intensities on longitudinal or cross-sections of tissue, with ultra-high resolution and accuracy of ion peaks (Dong et al. 2020; Nakabayashi et al. 2017; 2019b; Nakabayashi et al. 2020; Yamamoto et al. 2019). IMS analysis enables us to spatially acquire targeted or untargeted metabolome data. The resolution and accuracy of this analysis makes it an extremely powerful method to determine exactly where target metabolites accumulate.

Asparaptine is a sulfur (S)-containing metabolite found in green asparagus (*Asparagus officinalis*). Note that asparaptine shows inhibitory activity against angiotensin-converting enzyme, and it has been suggested that this metabolite may be effective for lowering blood pressure in animals, including humans (Nakabayashi et al. 2015; Nakabayashi and Saito 2017; Nakabayashi et al. 2019a). The biosynthesis of asparaptine is possibly derived from arginine and asparagusic acid, which is likely biosynthesized from valine (Mitchell and Waring 2014); however, the gene(s) controlling the asparaptine biosynthetic pathway are as yet unknown. As the initial step in identifying gene-to-metabolite associations in the biosynthesis of asparaptine is to understand the spatial control of asparaptine production, the accumulation and localization of metabolites within asparagus tissues need to be characterized.

In the present study, we performed MALDI-FTICR-IMS analysis to determine the localization of asparaptine in green asparagus spears. Segmentation analysis characterized tissue-redundancy of metabolites detected in asparagus spear sections. The results showed that asparaptine accumulates at high levels around developing lateral shoot tissues. The IMS results were validated by quantification of asparaptine in the lateral shoot using liquid chromatography-tandem mass spectrometry (LC-MS/MS). These results provide valuable information toward the regional function of asparaptine in asparagus, and suggest the lateral shoot tissues as a site of interest in gene-to-metabolite studies.

## Experimental section

### Plant materials

Commercial green asparagus products were analyzed using the IMS analysis. In this study, the result from the product of Niigata prefecture, which was purchased in a supermarket in Japan, was shown for the analysis of cross section. For the analysis of longitudinal sections, asparagus was harvested from Medicinal Plant Garden in Hoshi University.

### Preparing sections

Sections for IMS analysis and light microscopy were prepared as described in previous studies (Nakabayashi et al. 2017; 2019b). Briefly, for the IMS analysis, asparagus was cut with a razor, embedded with a frozen section compound (Surgipath FSC22: Leica Microsystems, Germany) and frozen in a −75 °C acetone bath (Histo-Tek Pino: Sakura Finetek Japan Co., Ltd., Tokyo, Japan). The frozen sample block was placed on a cryostat specimen disk and was cut with the knife blade until the desired tissue surface appeared. Adhesive film (Cryofilm Type II C (9), Section lab, Hiroshima, Japan) (Kawamoto 2003) was placed on the face of the block to obtain sections (each with a thickness of 20 μm) in the CM3050S cryostat (Leica Microsystems, Germany). The section on the glass slide was freeze-dried overnight at −30°C in the cryostat. The freeze-dried section on the film was attached to a glass slide (ITO coating, Bruker Daltonik GmbH) with cellophane tape. For light microscopy, frozen sections were stained with 0.05% toluidine-blue O solution for a minute and then washed with distilled water.

### MALDI-FTICR-IMS analysis

A 2,5-dihydroxybenzoic acid (DHB) matrix solution (30 mg/ml 80% MeOH including 0.2% TFA) was sprayed on the prepared section that was on the glass slide using ImagePrep (Bruker Daltonik GmbH) set to the default parameters. The freeze-dried section with the matrix was analyzed in the FTICR-MS SolariX 7.0 T instrument. The MALDI parameters are as follows: geometry, MTP 384 ground steel; plate offset, 100.0 V; deflector plate, 200.0 V; laser power, 50.0%; laser shots, 25; frequency, 2000 Hz; laser focus, medium; raster width, 60 μm. The MS conditions were as follows: mass range, *m/z* 100.32-400.00; average scan, 1; accumulation, 0.100 s; polarity, positive; source quench, on; API high voltage, on; resolving power, 66,000 at 400 *m/z*; transient length, 0.4893 s; Mode (data storage: save reduced profile spectrum, on; reduced profile spectrum peak list, on; data reduction, 95%; auto calibration: online calibration, on; mode, single; reference mass, *m/z* 273.039364); API Source (API source: source, ESI; capillary, 4500 V, end plate offset, −500; source gas tune: nebulizer, 1.0 bar; dry gas, 2.0 l/min; dry temperature, 180 °C); Ion Transfer (source optics: capillary exit, 220 V; detector plate, 200.0 V; funnel 1, 150.0 V; skimmer 1, 20.0 V; funnel RF amplitude, 180.0 Vpp; octopole: frequency, 5 MHz; RF amplitude, 350 Vpp; quadrupole: Q1 mass, 100.0 *m/z;* collision cell: collision voltage, −3.0 V; DC extract bias, 0.1 V; RF frequency, 1.4 MHz; collision RF amplitude, 1000.0 Vpp; transfer optics: time of flight, 0.500 ms; frequency, 4 MHz; RF amplitude, 350.0 Vpp); Analyzer (infinity cell: transfer exit lens, −20.0 V; analyzer entrance, −10.0 V; side kick, 5.0 V; side kick offset, −1.5 V; front trap plate, 0.540 V; back trap plate, 0.580 V; sweep excitation power, 12.0%; multiple cell accumulations: ICR cell fills, 1).

### Visualizing IMS data

Visualization was performed using flexImaging 4.2 (Bruker Daltonik GmbH, Bremen, Germany) using the IMS data. The data was normalized using Root Mean Square.

### Segmentation analysis

This analysis was performed using SCiLS Lab 2019c Core software. The IMS data was processed using the bisecting *k*-means method with the default parameters.

### Quantification of asparaptine

Asparaptine was quantified according to the previous study (Nakabayashi et al. 2015).

### Results and discussion

A cross-section of a green asparagus spear was scanned to determine the distribution of tissue types (**Figure 1A**). Cells were thickened around the shoot and lateral shoot tissues. Spatial metabolome data (41,353 spectra) were comprehensively acquired across the section in the range *m/z* 100–400 using MALDI-FTICR-IMS (**Figure 1B**). Segmentation analysis using the IMS data showed that the detected metabolites can be characterized into several patterns of signal intensities (**Figure 1C**). Six distinct patterns of signal intensities were obtained: each color meant a pattern that signal intensities at the parts are higher than other region. Except for the pattern colored as red, which meant non-section, six patterns were obtained. The analysis visualized the tissue-dependent and independent patterns. The orange, purple, green, and blue indicated the tissue dependency of the metabolites, whereas the light green and yellow indicated the independency that cannot be distinguished with the shape of the tissues. These patterns all implied that metabolites can play a local role within a region. Ion of asparaptine at *m/z* 307.08909 ([M + H]^+^; calculated for C_10_H_19_N_4_O_3_S_2_, 307.08930) was detected at a high level around the lateral shoot tissue (**Figure 1D**).

**Figure 1.**
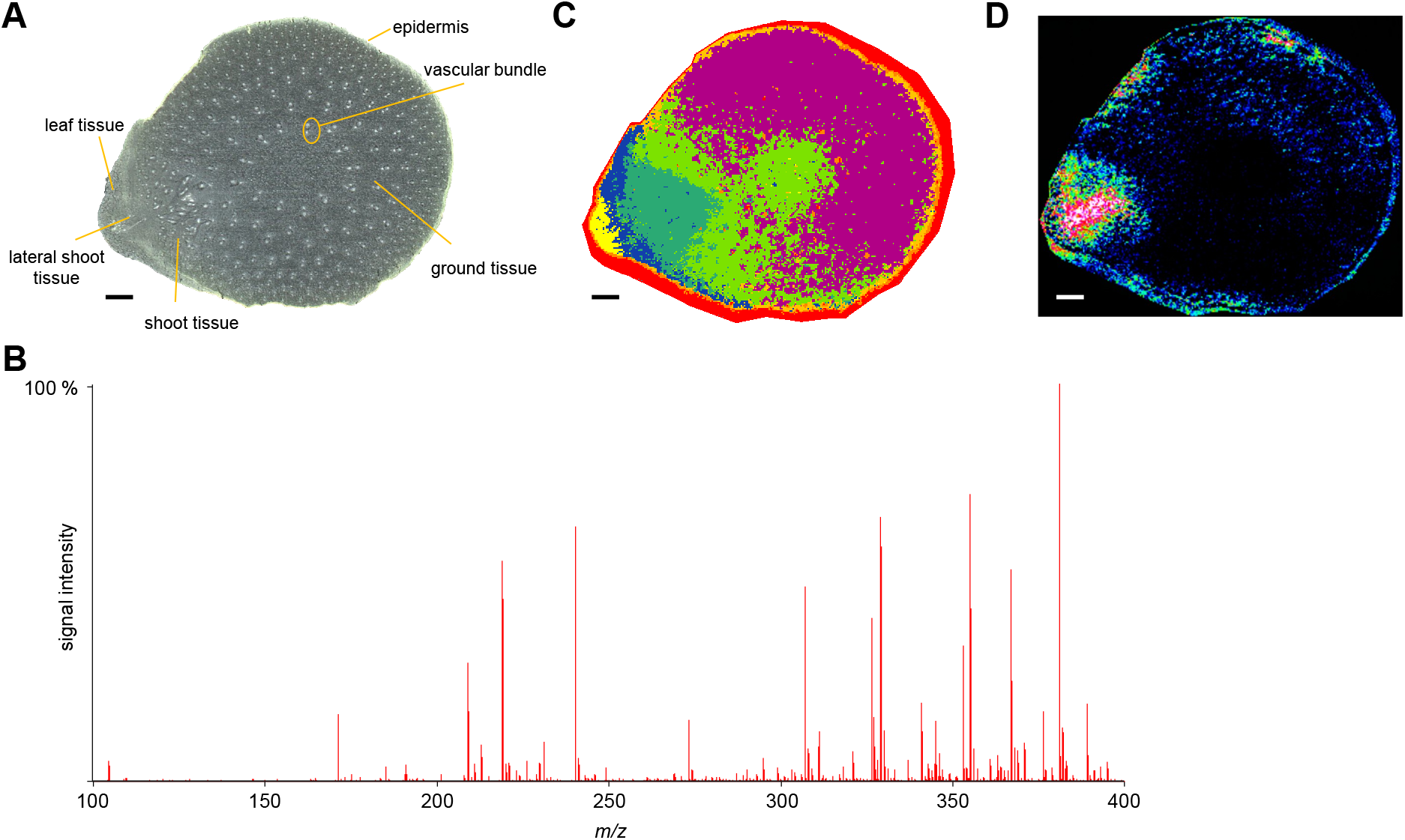
Spatial metabolomics in asparagus using matrix-assisted laser desorption/ionization Fourier transform ion cyclotron resonance imaging mass spectrometry (MALDI-FTICR-IMS). **A.** Scanning cross-section of an asparagus spear. **B.** Spatial metabolome data acquired in the range *m/z* 100–400. **C.** Segmentation analysis using the IMS data. Metabolites characterized into signal intensity patterns are indicated by color. **D.** Visualization of signal intensity at *m/z* 307.08909 ± 0.001 for asparaptine. Scale bar indicates 2 mm in panels **A, C**, and **D**.

To understand the characteristics of asparaptine in developing lateral shoot tissue, IMS analysis was performed on longitudinal sections prepared from green asparagus stems containing both developing and non-developing lateral shoots. We observed that asparaptine accumulates at high levels at the bottom of the shoot and increases with development of the lateral shoot (**Figure 2**).

**Figure 2.**
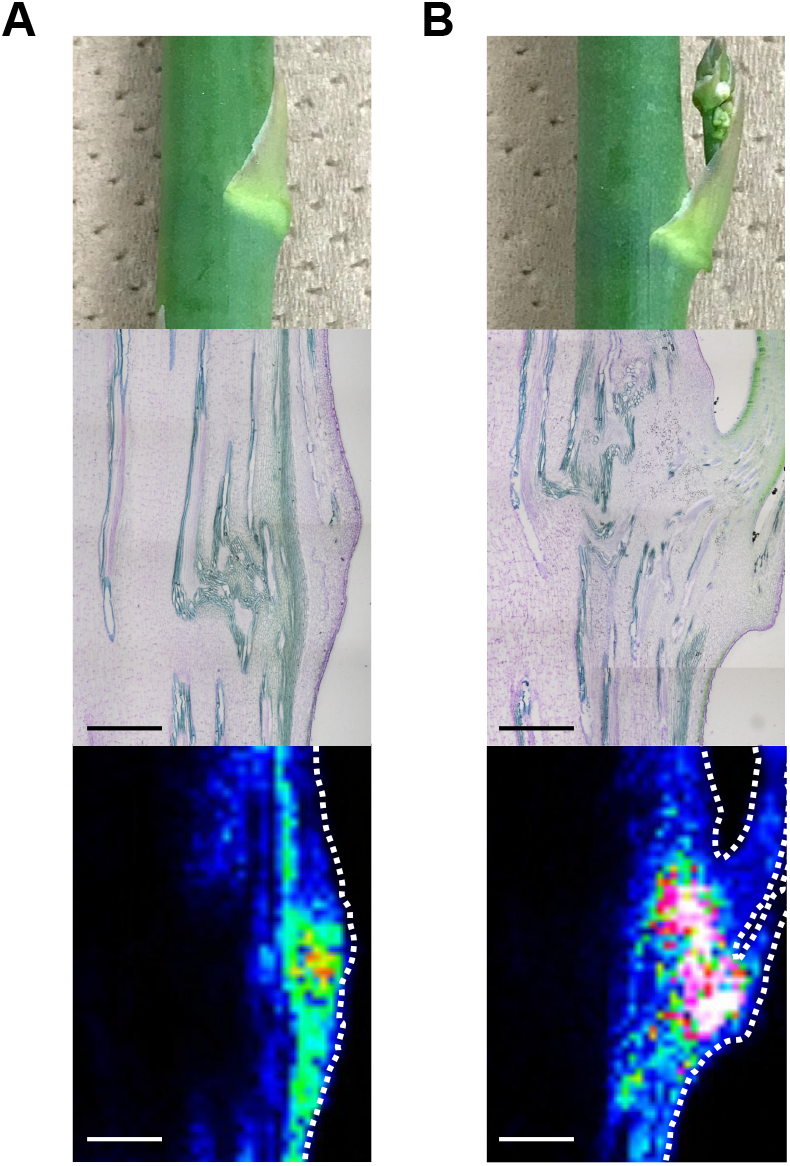
The behavior of asparaptine in the development of lateral shoots. Matrix-assisted laser desorption/ionization Fourier transform ion cyclotron resonance imaging mass spectrometry (MALDI-FTICR-IMS). MALDI-FTICR-IMS analysis of asparagus stem before (**A**) and after developing lateral shoots (**B**). Top panels show stems used in this study; middle panels show light microscopy images of longitudinal sections; bottom panels show IMS analysis of asparaptine. Black indicates lower level detected; white indicates higher level detected. Scale bar indicates 2 mm.

To confirm the localization of asparaptine in lateral shoot tissues, the lateral shoots were dissected away from the spears, and the amount of asparaptine in the lateral shoots and remaining spears was quantified against an asparaptine standard curve using LC-MS/MS. The amount of asparaptine was higher in lateral shoots than in the remaining tissue, supporting the results obtained using the IMS analysis (**Figure 3A**). As asparagus is a perennial plant, the stems wither during winter in Japan. In an LC-MS/MS analysis of asparagus stems collected from March through to the subsequent January, we observed a seasonal change in asparaptine levels, suggesting that asparaptine accumulates in actively growing tissues (**Figure 3B**).

**Figure 3.**
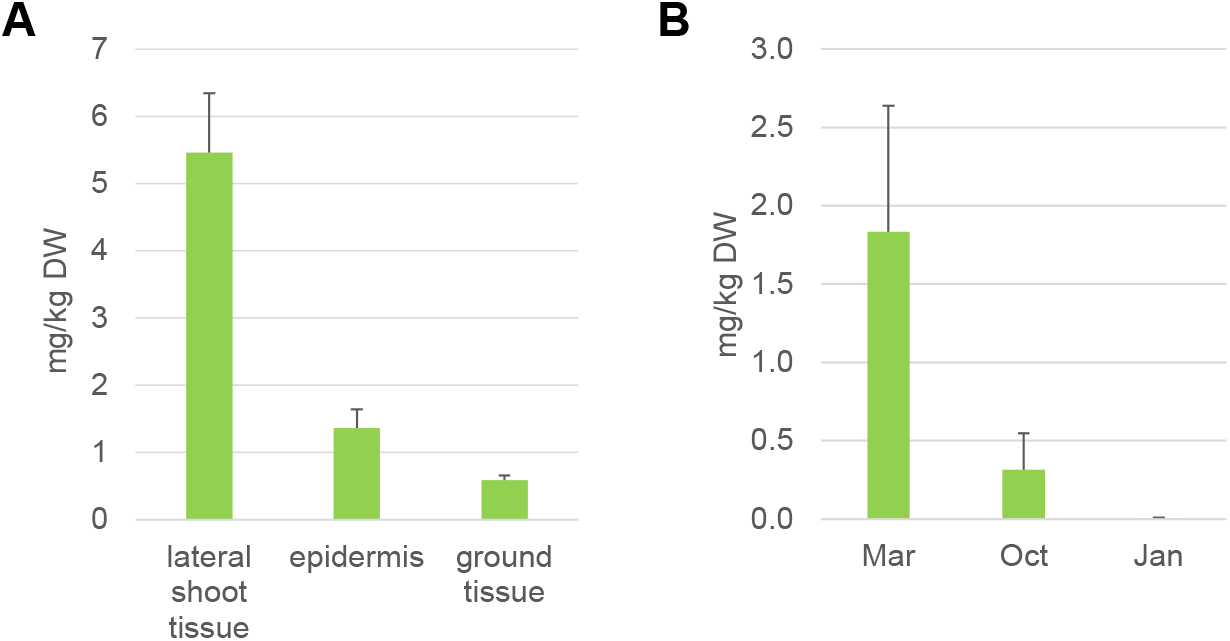
Quantification of asparaptine using liquid chromatography-tandem mass spectrometry (LC-MS/MS). **A.** Quantification of the amount of asparaptine in spears. The spears were harvested in March 2016. **B.** Seasonal variation in asparaptine level. Spears (March 2015), withering stems (October 2015), and withered stems (January 2016) were harvested and analyzed. The samples all were harvested in Medicinal Plant Garden of Hoshi University.

To investigate the possibility that asparaptine biosynthesis occurs at the lateral shoot tissue, the IMS data of the cross-section were employed to visualize the regional signal intensity of the arginine ion at *m/z* 175.11871 ([M + H]^+^; calculated for C_6_H_15_N_4_O_2_, 175.11895) ± 0.001 (**Figure 4**). This visualization suggests that arginine is present in the lateral root, where we previously observed asparaptine distribution; however, arginine is also detected more widely outside the lateral root. The IMS analysis could not detect ions of asparagusic acid or valine. The parameters of the instrument or the matrix reagent used in this study need to be changed to improve this.

**Figure 4.**
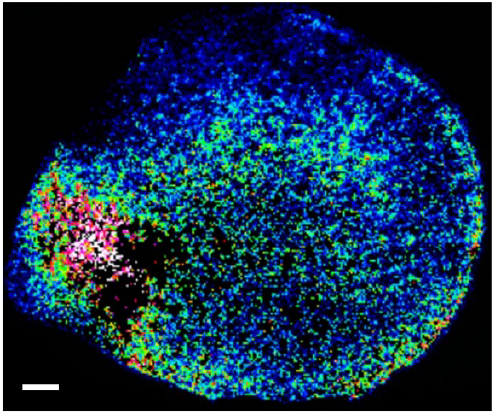
Visualization of arginine using matrix-assisted laser desorption/ionization Fourier transform ion cyclotron resonance imaging mass spectrometry (MALDI-FTICR-IMS). The signal intensities for arginine at *m/z* 175.1187 ± 0.001 were visualized in asparagus. Black indicates lower level detected; white indicates higher level detected. Scale bar indicates 2 mm.

As described above, the biosynthetic pathway for asparaptine is as yet unknown but is thought to involve conjugation with arginine. In the present study, we observed differential localization of asparaptine and arginine, with asparaptine mainly localized in the lateral root tissues, whereas arginine was more widely localized in the tissue section. These data strongly suggest that the reaction for asparaptine biosynthesis through conjugation with arginine occurs at the lateral shoot tissues. Therefore, it is likely that the biosynthetic gene(s) involved in the conjugation are mainly expressed in the lateral shoot tissue. These results provide a solid foundation for performing local multiomics studies to characterize the expression of candidate genes in the regions of high and low asparaptine accumulation. Co-behavior analysis to reveal genes with an expression pattern mimicking the pattern of asparaptine accumulation will enable the identification of gene(s) controlling the asparaptine biosynthetic pathway.

In summary, we performed spatial metabolomics using MALDI-FTICR-IMS to identify the localization of asparaptine in cross- and longitudinal sections of asparagus. The IMS analysis showed that asparaptine accumulates at high levels in lateral shoot tissue, together with arginine. The localization of asparaptine at the lateral shoot invites the possibility of local multiomics for identifying the gene(s) involved in its biosynthetic pathway. Once these biosynthetic genes are identified, the function of asparaptine may be determined using genome-editing techniques. The spatial metabolomics study here using the IMS analysis also characterized the localization of other metabolites in addition to asparaptine. The pattern of metabolite distribution suggests that metabolites have local functions within asparagus tissues. The functions of most metabolites, especially specialized metabolites, are largely unknown. The distribution patterns of the metabolites detected in this study may provide clues to their specific function within particular regions of asparagus spear tissue. Spatial metabolomics is a crucial tool in the identification of metabolite function.

## Acknowledgements

We would like to thank Yoshihiko Morishita and Takashi Nirasawa (Bruker K. K.) for advices on the IMS analysis.

## Author Contributions

R.N. designed the research. R.N. and H. S. harvested the asparagus samples. R.N., K. H., and K. T. prepared the sections. R. N. performed the IMS analysis and analyzed the data. T. M. performed the quantification. All the authors discussed the research. R. N. wrote the manuscript.

## Notes

The authors declare no competing financial interest.

